# GLADE: Accurate inference of Gains, Losses, Ancestral genomes, and Duplication Events for comparative genomics

**DOI:** 10.64898/2026.01.27.702036

**Authors:** Laurence J. Belcher, Steven Kelly

**Author notes:** Steven B. Kelly **Email:**. **Author Contributions:** LJB and SK designed research; LJB performed research and analyzed data; LJB and SBK wrote the paper. Competing Interest Statement: SK is co-founder of Wild Bioscience Ltd and an employee of Ellison Institute of Technology, Oxford Limited.

## Abstract

Changes in gene content through gain and loss play a key role in the adaptation and diversification of species. Accordingly, our ability to detect and accurately document the history of these changes is important for our understanding the evolutionary trajectories of life on Earth. Here we present GLADE, a tool that accurately reconstructs gene gains, losses, and duplications for a set of species under consideration and uses this information to infer ancestral gene contents for every speciation event in the species tree. GLADE requires as input only a standard OrthoFinder results directory, and outputs the full evolutionary history of every orthogroup, including branch-specific changes and reconstructed ancestral genomes. We benchmark GLADE using both real and simulated data and show that GLADE accurately identifies orthogroup gains, losses, and duplications, and reconstructs ancestral orthogroup sizes with higher precision and overall accuracy than any competitor method. To illustrate the utility of the method, we apply GLADE to a dataset of 78 mammalian genomes and uncover repeated contractions in orthogroups associated with tooth formation on branches leading to ant- and termite-eating mammals - revealing convergent genomic signatures underlying this dietary specialization. GLADE and accompanying documentation and tutorials are freely available at https://github.com/lauriebelch/GLADE/.

## Introduction

Every species’ genome has been shaped by a continual churn of gene gain, loss, and duplication over evolutionary time. This genomic churn plays a key role in the adaptation and diversification of species across all domains of life. For example, expansions in select sets of orthogroups contributed to plants colonising terrestrial environments (Leebens-Mack et al. 2019; Bowles et al. 2020) and animals emerging from their simple ancestor to develop complex multicellularity (Paps and Holland 2018; Ocaña-Pallarès et al. 2022). Similarly, contractions in specific orthogroups allowed fungi to diversify into new niches (Merényi et al. 2023; Feng et al. 2025), and enabled symbionts adapt to a sheltered life within their hosts by convergently losing redundant genes (Kohler et al. 2015; Kelly 2021; Siozios et al. 2024). In addition to altering individual adaptive processes, these genomic changes can also feedback in co-evolutionary relationships. For example, as leaf-cutter ants began farming fungi, contractions in genes encoding digestive enzymes were matched by reciprocal expansions in their fungal partners (Nygaard et al. 2016). Similarly, in bacteria, patterns of gene gain, loss, and duplication are crucial in the evolution of virulence and pathogenicity (Hall et al. 2017; Murray et al. 2021) and shaped by variation in lifestyle (Brockhurst et al. 2019; Dewar et al. 2024). Thus, changes in gene content are associated with adaptation across the tree of life.

To enable the discovery and analysis of changes in gene content over evolutionary time, it is important to know the phylogenetic history of genes (Fitch 1970; Linard et al. 2021). This is important as it enables us to distinguish precisely which members of an orthogroup were lost or duplicated during evolution, and thus enable an accurate understanding of the evolutionary dynamics of genes and genomes (Koonin 2005). This accurate reconstruction cannot be obtained by count data alone, and it is poorly resolved by simple sequence similarity approaches (Gabaldón 2008; Emms and Kelly 2015). With the advances in sequencing technology and the continued increase in the availability of genomic information (Lewin et al. 2018; The Darwin Tree of Life Project Consortium 2022) there is a real need for methods that can analyse diverse datasets to chart the changes in gene content that shaped extant genomes, and enable us to infer the changes that are associated with adaptation and diversification (Sarton-Lohéac et al. 2025).

Currently, the most widely used method for interring changes in gene content along a phylogeny is CAFÉ (Mendes et al. 2021). Given a species tree and a file with orthogroup counts in extant species, CAFÉ fits a birth-death model to estimate the changes in gene content that best explain the distribution of orthogroup sizes we observe in living species (Hahn et al. 2005; De Bie et al. 2006). Users can specify how many different categories of evolutionary rates the model should use, allowing some orthogroups to evolve slower or faster than others. Although CAFÉ is widely used, the method has a key limitation. CAFÉ only uses count data and is thus unable to resolve which paralog within an orthogroup was lost or duplicated at a given node in the species tree. This limits the information that can be gleaned about the evolutionary history of orthogroups, and can lead to mistaken inferences about which genes have been duplicated or lost. Thus, there is a requirement for methods that reconstruct ancestral changes in gene content that are accurate, defined by phylogeny, and scalable to contemporary comparative genomic research.

Here, we present GLADE, an automated and easy to use tool that provides phylogenetic reconstruction of gene gain, loss, and duplication within sets of orthogroups. GLADE only requires an OrthoFinder (Emms and Kelly 2015; Emms and Kelly 2019; Emms et al. 2025) results directory as input, and uses the gene trees, species tree, and orthogroup information therein to infer and analyse the complete set of gains, losses, and duplications within each orthogroup, mapping every event to its precise location on the species tree. GLADE then uses this information to reconstruct ancestral proteomes for every node in the species tree. We show that GLADE can accurately reconstruct these evolutionary events and demonstrate its utility in identifying convergent changes in orthogroup sizes during the convergent evolution of myrmecophagy within a dataset of 23,000 orthogroups across 78 mammals.

## Results

### GLADE Algorithm overview

The GLADE algorithm is described in detail in the ‘Methods’ section and described briefly here. GLADE takes as input the standard output files produced by OrthoFinder v3, including a species tree, a file containing orthogroups, and a gene tree for each orthogroup.

GLADE first performs two major steps for each orthogroup: 1) Identification of the node on the species tree where the orthogroup emerged. 2) Identification of subsequent loss and duplication events in the gene tree for the orthogroup. All events are then mapped to the branch or node on the species tree where they occur. This allows GLADE to infer the size of each orthogroup at every internal node in the species tree. The tool then builds a reconstructed pseudo-genome FASTA file for each node in the species tree (Figure 1).

**Figure 1:**
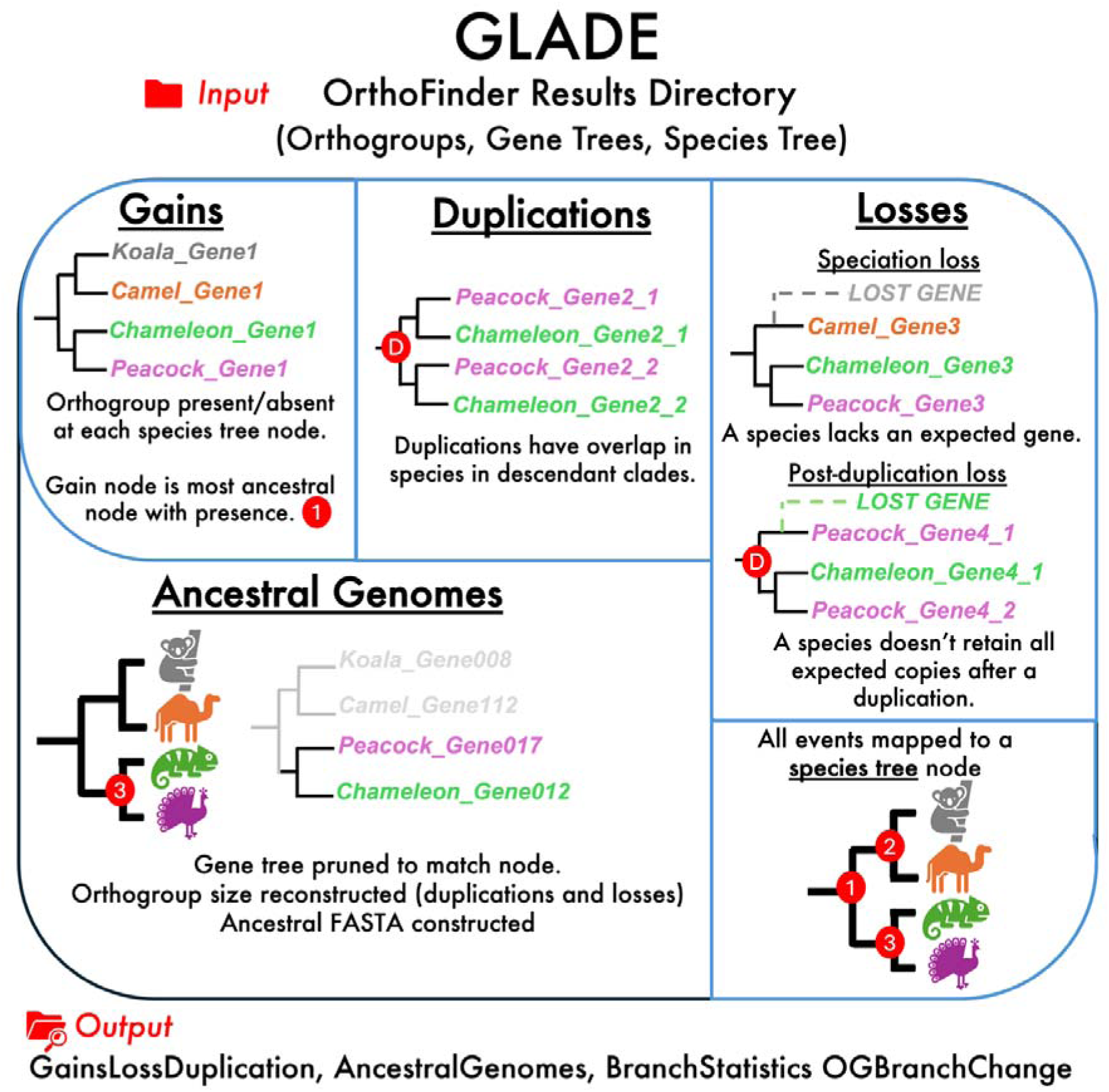
Overview of the GLADE workflow. GLADE takes as input the results directory from OrthoFinder (reading the orthogroups, gene trees, and species tree files) and identifies evolutionary events by reconciling gene trees to the species tree. Gains are inferred as the most ancestral nodes where an orthogroup is present. Duplications are detected when gene copies from the same species overlap in descendant clades, while losses are identified when expected genes are missing after speciation or duplication events. All events are summarized and mapped to the species tree, producing outputs detailing all evolutionary events, the relevant genes and orthogroups, and mappings from gene to species trees. From these mappings, GLADE reconstructs ancestral gene content and generates ancestral genome FASTA files for each internal node in the species tree.

### Simulations can produce realistic orthogroups for benchmarking

To evaluate the performance of GLADE requires a dataset where the complete evolutionary history of a set of orthogroups in a set of species is fully known. This necessitates resolved phylogenetic trees for a full set of orthogroups with all gene duplications, gains and losses annotated and reconciled with the species tree. No such benchmark dataset exists. For example, the expert curated OrthoBench dataset only provides a set of phylogenetic trees for 70 reference orthogroups across 12 species (Trachana et al. 2011; Emms and Kelly 2020). This dataset represents <1% of all genes in those species and thus is a limited sample size for comparative testing and benchmarking. Moreover, these trees are also only inferences, whilst the true phylogenetic trees are not known. Other resources such as Ancestral Genomes (Huang et al. 2019) and OMA (Altenhoff et al. 2024) provide ancestral genome reconstructions based on hierarchical orthogroups, but the absence of a known ground truth again precludes objective benchmarking against these approaches. As suitable benchmark datasets were not available, we instead sought to simulate datasets whose properties matched as close as possible to those of real datasets.

To generate these simulations, we used two empirical datasets – one containing 100 species of budding yeast (Shen et al. 2018), and the other with 50 species of plants (Bolser et al. 2016). To generate sets orthogroups for which parameters for the simulation could be trained, the proteomes were subject to orthogroup inference using Broccoli (Derelle et al. 2020), SonicParanoid2 (Cosentino et al. 2024), FastOMA (Majidian et al. 2025), and OrthoFinder (Emms et al. 2025). Multiple disparate methods were used here to prevent biasing the simulations towards any particular orthogroup inference method. The orthogroups produced by these methods were then subject to phylogenetic tree inference and the properties of the orthogroups and the resultant orthogroup trees were analysed used to train the parameters of the corresponding simulations. Once determined, the trained parameters were then used to simulate a set of orthogroups, with each orthogroup seeded by a real gene sampled at random from a starting proteome.

Simulations were produced using a custom multi-step method (see Methods for full details). For each orthogroup, a random gene was selected from the real starting genome. SaGePhy (Kundu and Bansal 2019) was then used to simulate a gene tree within the real species tree, allowing for duplication, losses, and variable rates of evolution along different branches. IQ-TREE’s AliSim module (Ly-Trong et al. 2022; Wong et al. 2025) was then used to simulate a multiple sequence alignment corresponding to the gene tree, allowing for different rates of sequence evolution in different regions of the protein (see Methods). The resultant simulated sequences from each orthogroup were then distributed to their relevant species proteome. Multiple rounds of parameter training were performed until the simulated trees and orthogroup sizes exhibited properties that matched those observed for the training datasets.

For the fungi dataset we simulated 5,917 orthogroups from the *Saccharomyces cerevisiae* genome (Engel et al. 2013). For the plant dataset we simulated 27,000 orthogroups from the *Arabidopsis thaliana* genome (The Arabidopsis Genome Initiative 2000). The final set of simulated orthogroups and gene trees were then analysed and compared to those observed in the real-world training data (Figure 2). This revealed that the simulated orthogroups had analogous distributions of the number of species and genes per orthogroup to those obtained real data (Figure 2 A&B, E&F). We also compared the phylogenetic trees produced by simulation to those in the real datasets using two different tree metrics that capture overall tree topology and structure. Again, the distribution of Wiener indices, the sum of all pairwise path lengths, is a close match between simulated and empirical orthogroups (Figure 2 C&G), demonstrating that the simulated gene trees recreate realistic patterns in trees. Similarly, the treeness index, which is the proportion of total branch length contained in internal (non-terminal) branches, is highly similar between simulated and real data (Figure 2 D&H). This indicates that the relative concentration of evolutionary change along internal versus terminal branches is consistent with the empirically observed values from real gene trees. Importantly, these values are similar across all orthology inference tools and is thus not an inherent property of any particular method. It is also important to note that whilst several parameters of the simulation were estimated directly from empirical data (e.g. the gamma shape alpha, the proportion of invariant sites, and the duplication rates), the metrics in Figure 2 demonstrate emergent properties of the simulated orthogroups which were not directly modelled.

**Figure 2:**
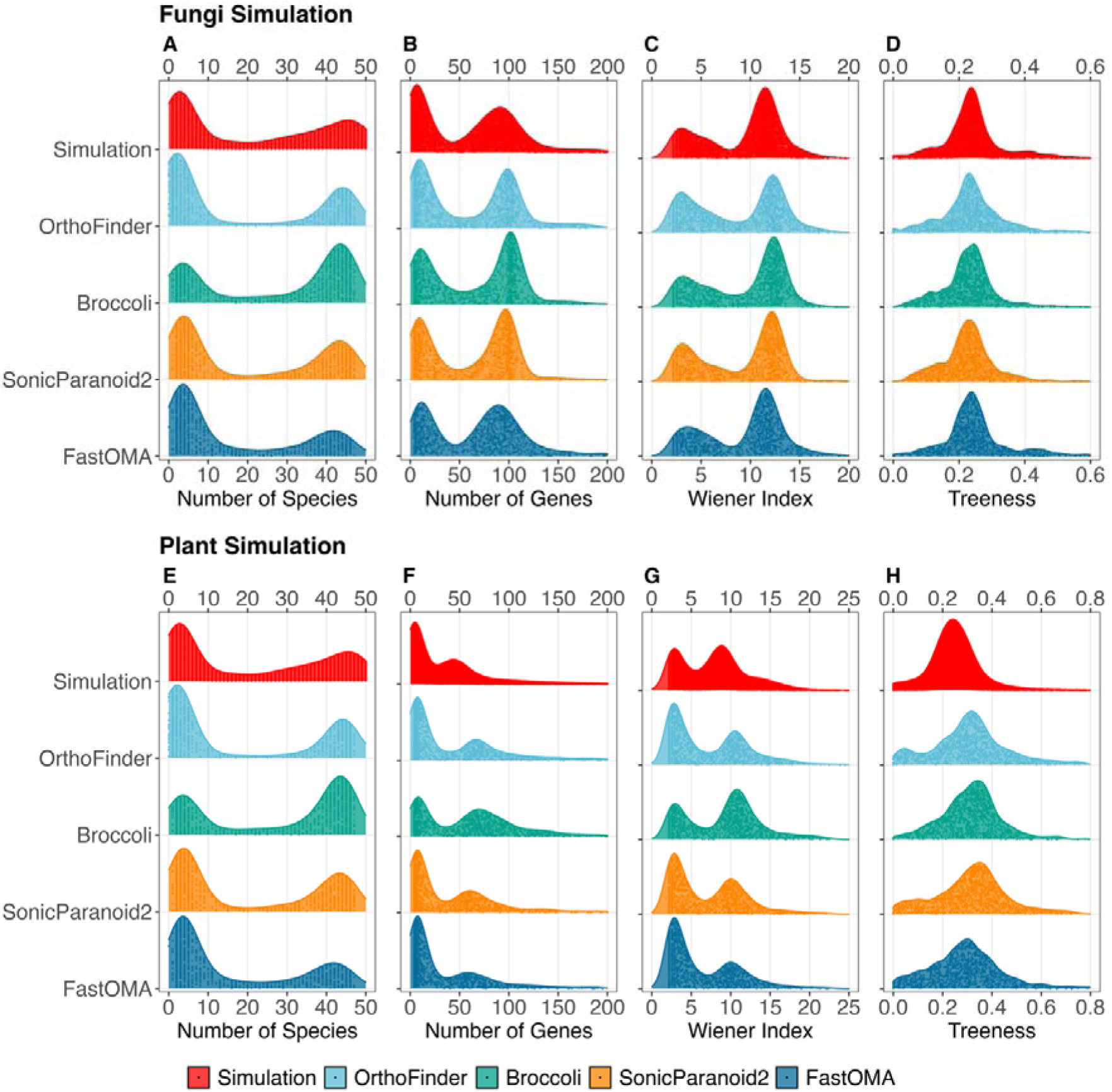
Comparison of simulated orthogroups to real orthogroups inferred by multiple methods. The top row (Panels A-D) show data from the fungi simulation. The bottom row (Panels E-H) show data from the plant simulation. (A&E) show number of species present in each orthogroup. (B&F) show number of genes present in each orthogroup. (C&G) shows Wiener index of the gene trees inferred from each orthogroup. (D&H) show Treeness of the gene trees inferred for each orthogroup. Distributions for the simulated orthogroups are compared to those obtained by orthogroup inference using OrthoFinder, Broccoli, SonicParanoid2, and FastOMA.

As the simulated datasets contain a completed set of resolved gene trees with all gene, gains, losses and duplications fully resolved against the species tree, this enables us to benchmark the performance of GLADE to competitor methods. The two simulations also had different properties, allowing us to test the accuracy of GLADE under different evolutionary scenarios.

### GLADE accurately reconstructs gains, losses, and duplications

To evaluate the accuracy of GLADE in reconstructing the evolutionary history of orthogroups, we compared its performance on our simulated dataset to an alternative method FastOMA, which also provides data on events in the evolution of an orthogroups (extractable from the FastOMA output using pyHAM (Train et al. 2019)). Whilst other tools can perform individual components of what GLADE does (e.g. identifying duplications on a gene tree), FastOMA is the only comparator method that allows a comparable reconstruction of orthogroup history and is therefore the most appropriate comparison. For orthogroup gain, gene loss, and gene duplication events, we extracted the ground truth from our simulation output, and quantified precision, recall, and F1 score by comparing true events to those predicted by both methods. For this, we use a weighting approach that accounts for non-perfect overlap between predicted orthogroups and true orthogroups to avoid biasing the results towards false predicted orthogroups that show only marginal overlap with a reference orthogroup. We also measured the magnitude of errors made by each tool, to give a more complete view of accuracy that looks at how close errors are to the truth. For metrics that require placing events on the correct node of the species tree we measured errors in two ways – counting either number of nodes between truth and prediction (topological error), or summing branch length. We tested for differences in median error magnitudes between tools using permutation steps where tools labels are randomly reassigned to generate a null distribution where tools have equivalent errors.

### Orthogroup gain

GLADE showed higher accuracy than FastOMA in identifying orthogroup gains, defined as the node on the species tree at which an orthogroups first emerged (Figure 3). This is true for both simulations, with higher F1 scores in the fungi dataset (97.0 vs. 88.2), and the plant dataset (92.4 vs. 71.7). GLADE also achieved higher precision (fungi: 95.4 vs. 88.5; plant: 90.7 vs. 70.9), and higher recall (fungi: 98.6 vs. 88.0; plant: 94.1 vs. 72.5) (Figure 3A&C). In the fungi simulation, the two methods showed no difference in topological accuracy of gain node location, with both exhibiting a median error of two nodes (Figure 3B). By contrast, in the plant simulation GLADE made significantly smaller topological errors than FastOMA (median 2 vs. 3 nodes, permutation test p<10^-4^) (Figure 3E). In both simulations, GLADE also produced significantly smaller errors when error magnitude was quantified using branch length rather than topological distance (fungi: median 0.17 vs 0.24, p<10^-4^; plant: median 0.25 vs 0.37, p<10^-4^) (Figures 3C&F), indicating that GLADE achieved closer placement of predicted gain nodes to true nodes in terms of evolutionary distance.

**Figure 3:**
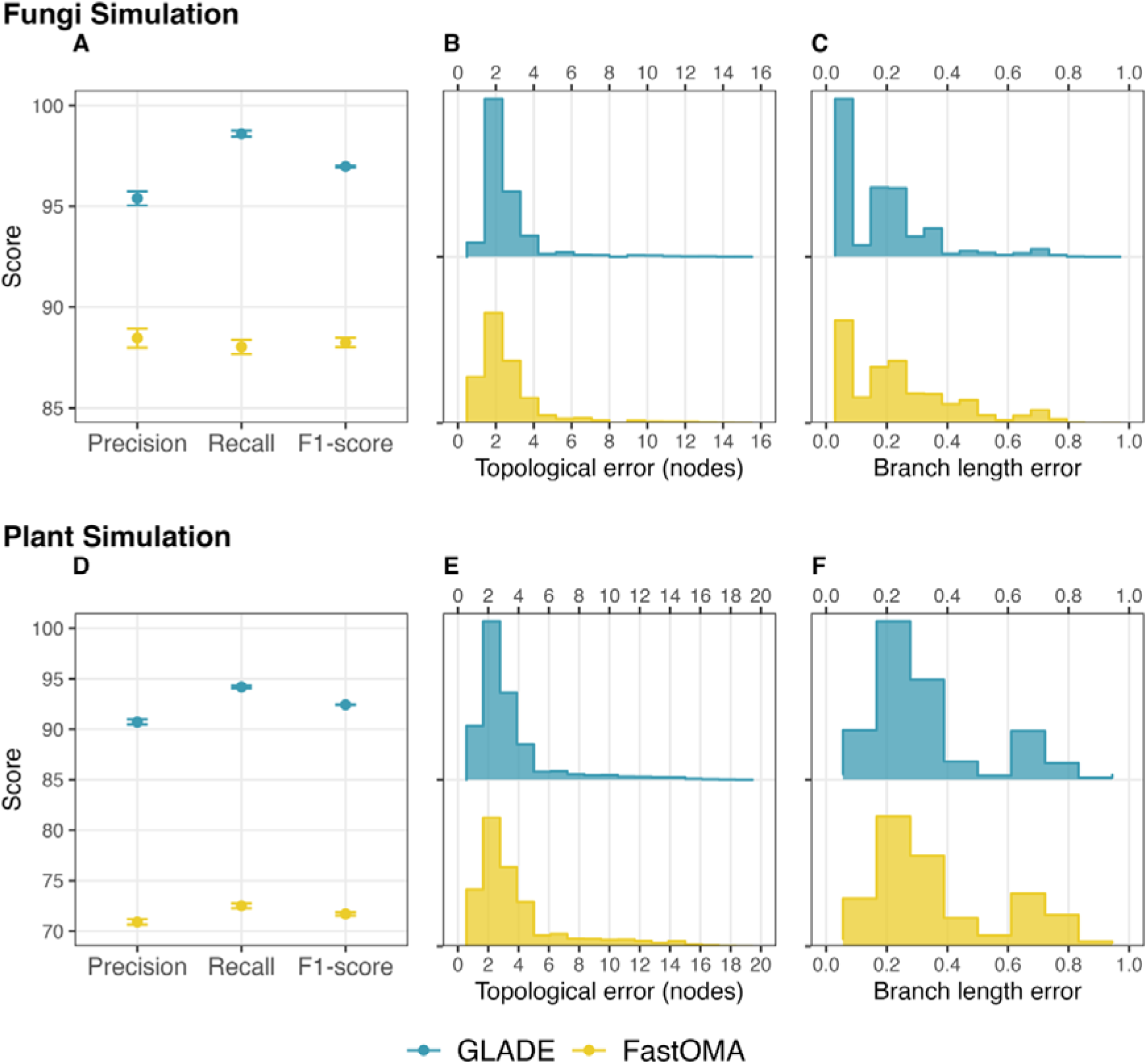
Benchmarking GLADE on orthogroup gain inference. The top row (Panels A-C) show data from the fungi simulation and the bottom row (Panels D-F) show data from the plant simulation. (A&D) show precision, recall, and F1-score for GLADE (blue) and FastOMA (yellow). (B&E) show the distribution of topological errors (number of nodes between true and predicted gain location). (C&F) show the distribution of branch length error (total branch length on path between true and predicted node). Error bars on panels A&D show standard error of the mean value across all true orthogroups.

### Gene loss

GLADE showed higher accuracy than FastOMA in placing gene loss events on the species tree (Figure 4). This higher accuracy was manifest in both simulations, with higher F1 scores in the fungi dataset (81.2 vs. 68.4), and the plant dataset (62.9 vs. 47.1). GLADE also achieved higher precision on both datasets (fungi: 85.6 vs. 60.3; plant: 63.6 vs. 38.8). FastOMA had 2% higher recall than GLADE on the fungi dataset (79.0 vs. 77.1), but this was countered by its 30% lower precision. On the plant dataset, GLADE scored higher recall than FastOMA (62.1 vs. 60.3) (Figure 4A&C).

**Figure 4:**
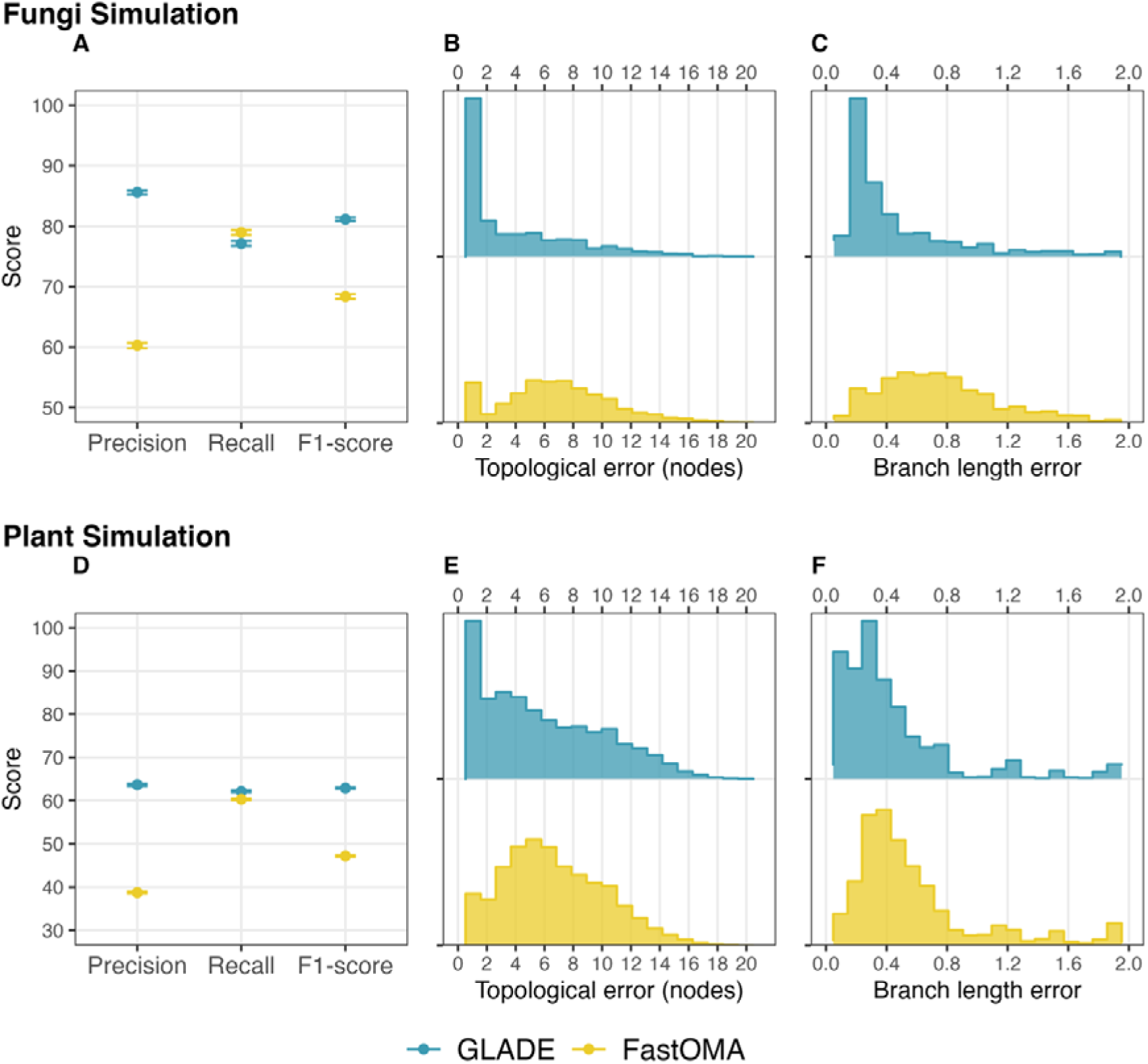
Benchmarking GLADE on gene loss inference. The top row (Panels A-C) show data from the fungi simulation and the bottom row (Panels D-F) show data from the plant simulation. (A&D) show precision, recall, and F1-score for GLADE (blue) and FastOMA (yellow). (B&E) show the distribution of topological errors (number of nodes between true and predicted loss location). (C&F) show the distribution of branch length error (total branch length on path between true and predicted node). Error bars on panels A&D show standard error of the mean value across all true orthogroups.

Where GLADE made errors, it also tended to make smaller errors than FastOMA on both datasets. In the fungi simulation, topological errors were on average almost five nodes smaller for GLADE compared to FastOMA (median 2 vs. 7, permutation test p<10^-4^). In the plant simulation where both tools were less accurate overall, GLADE made significantly smaller topological errors than FastOMA (median 5 vs. 6, permutation test p<10^-4^). In both simulations, GLADE also produced significantly smaller errors when error magnitude was quantified using branch length rather than topological distance (fungi: median 0.28 vs. 0.70, p<10^-4^; plant: median 0.33 vs. 0.45, p<10^-4^) (Figures 4C&F), indicating that GLADE was closer to the truth in loss events that it inferred incorrectly.

### Gene duplication events

GLADE showed higher accuracy than FastOMA in identifying gene duplication events. (Figure 5). A true positive occurs when a tool infers a correct set of leaves involved in a duplication event. False positives are predicted duplications that don’t exist in the ground truth, and false negatives are true duplications that aren’t inferred by a tool.

**Figure 5:**
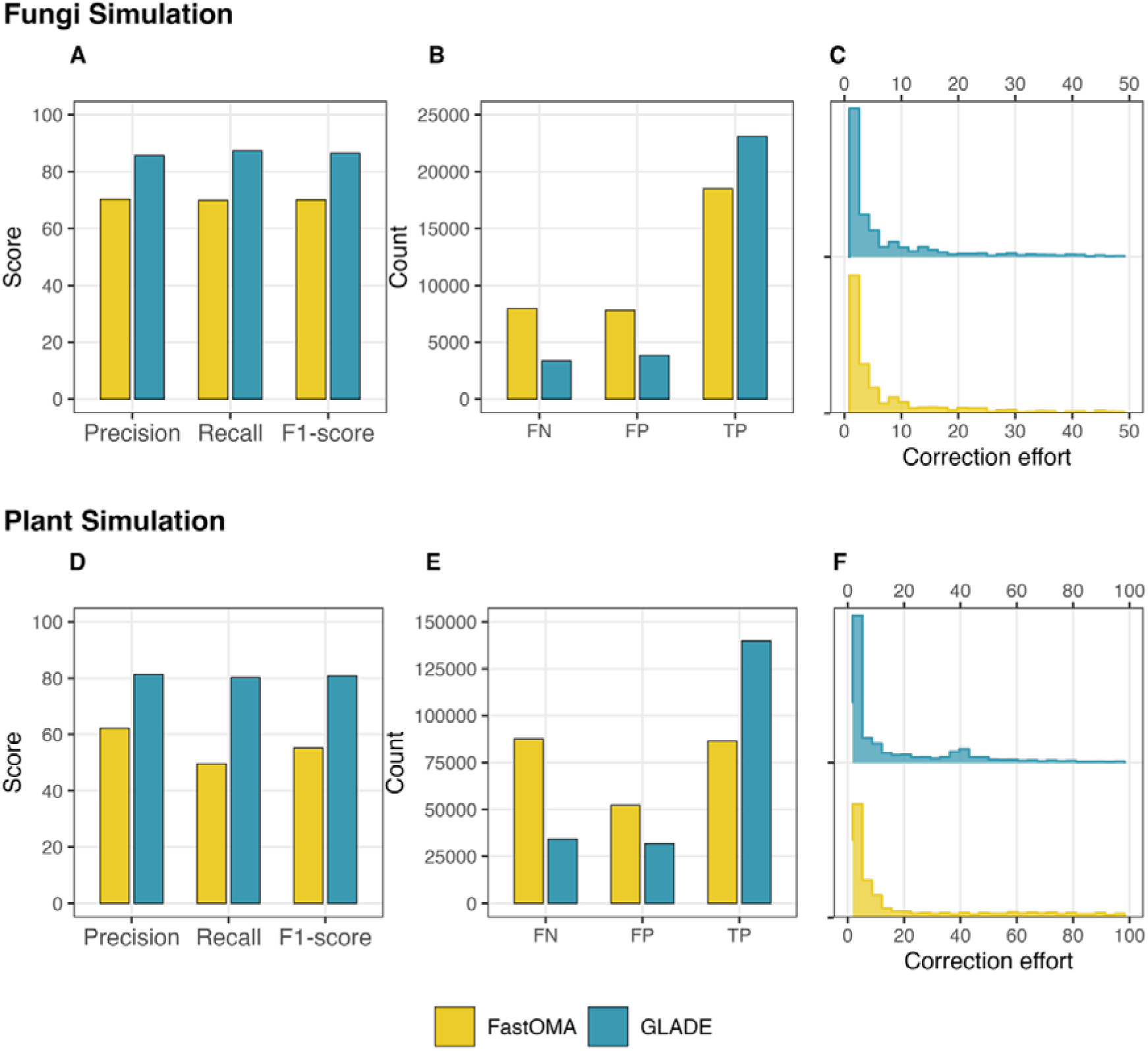
Benchmarking GLADE on gene duplication event inference. The top row (Panels A-C) show data from the fungi simulation and the bottom row (Panels D-F) show data from the plant simulation. (A&D) show precision, recall, and F1-score for GLADE (blue) and FastOMA (yellow). (B&E) show the counts of true positives, false positives, and false negatives for each tool. (C&F) show the distribution of correction effort, defined as the number of genes that must be added or removed to turn an incorrect gene duplication event into a correct one.

GLADE was more accurate than FastOMA in both simulations, with higher F1 scores in the fungi dataset (86.5 vs. 70.2), and the plant dataset (80.8 vs. 55.4). GLADE also achieved higher precision on both datasets (fungi: 85.7 vs. 70.4; plant: 81.4 vs. 62.3) and also higher recall (fungi: 87.3 vs. 70.0; plant: 80.3 vs. 49.7). A breakdown of these errors shows that GLADE doesn’t just find more true positives (fungi 23,094 vs. 18,497; plant 139,751 vs. 86,437), it also finds fewer false positives (fungi 3,863 vs. 7,795; plant 31,838 vs. 52,326), and fewer false negatives (fungi 3,362 vs. 7,959; plant 34,346 vs. 87,660) (Figure 5B&E).

We measured the size of errors by matching each incorrect duplication to its closest true duplication using Jaccard distance. The magnitude of the error was then quantified as the amount of ‘correction effort’ required to turn a false duplication into a true one, where adding or removing one gene is equivalent to one unit of effort. In the fungi simulation, there was no difference in magnitude of correction effort required to transform a false duplication into a true one. In contrast, in the plant simulation the required correction effort was higher for FastOMA than it was for GLADE (median effort 5 vs. 4, permutation test P<10^-4^), indicating that GLADE’s mistakes were of smaller magnitude (Figure 5C&F).

### GLADE accurately reconstructs orthogroup sizes at ancestral nodes

We next assessed the accuracy of ancestral orthogroup size reconstruction. For this comparison, we compared GLADE to three alternative methods for reconstructing gene family evolution – CAFÉ (Mendes et al. 2021), BadiRate (Librado et al. 2012), and ancestral character estimation (ace) (Revell 2014). CAFÉ is a highly cited method which uses a birth-death model to infer orthogroups size changes. BadiRate is a similar method which implements an extended innovation model that allows new orthogroups to emerge along the species tree. Ace uses maximum likelihood to estimate change in a trait’s state along a species tree under a model of discrete character evolution. Whilst not specifically designed for orthogroups, the general methodology is applicable to the problem of reconstructing ancestral orthogroup sizes. Along with GLADE itself, all methods require as input a species tree and a dataset showing orthogroup sizes in extant species at the tips.

GLADE was the most accurate method in both simulations, with the highest F1 score on both the fungi (94.9) and plant (86.8) simulations. Both CAFÉ and BadiRate score similar or greater recall than GLADE (fungi: CAFÉ=98.9, BadiRate=98.4, GLADE=98.6; plants CAFÉ=93.1, BadiRate=90.9, GLADE=92.5), however this is more than countered by GLADE’s superior precision (fungi: CAFÉ=86.3, BadiRate=84.9, GLADE=91.4, plants CAFÉ=70.3, BadiRate=72.2, GLADE=81.8).

Ace was the second-best performing method overall, achieving F1 scores of 94.8 on the fungi dataset and 84.1 on the plant dataset. All tools tend to make only small errors with a median error size of one (Figure 6B&D), and there were no significant pairwise differences in error magnitude between tools.

**Figure 6:**
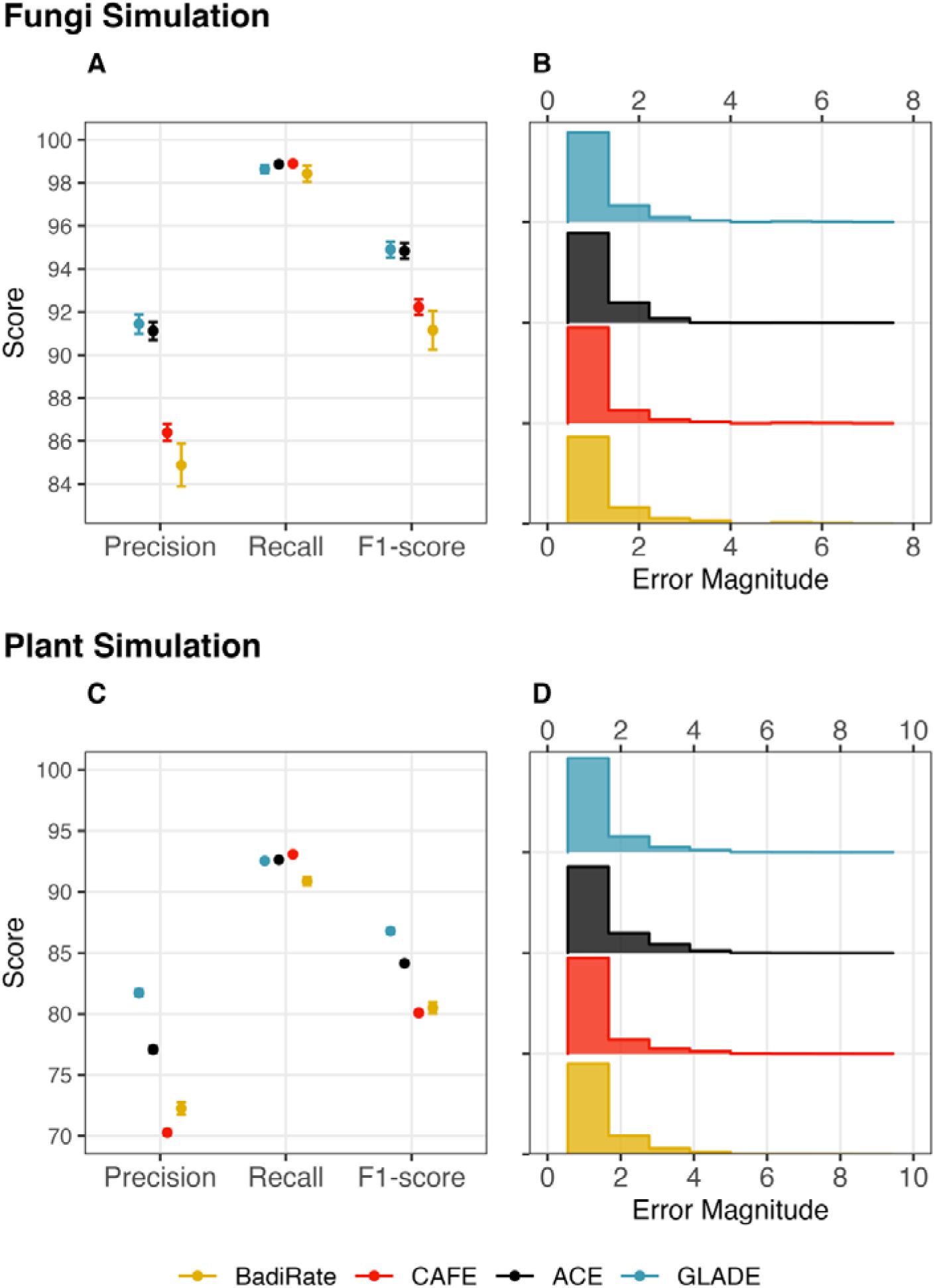
Benchmarking GLADE on ancestral orthogroups size reconstruction. The top row (Panels A&B) show data from the fungi simulation and the bottom row (Panels C&D) show data from the plant simulation. (A&C) show precision, recall, and F1-score for GLADE, ACE, BadiRate, and CAFÉ. (B&D) show the distribution of magnitude of error, defined as the absolute difference between inferred at predicted orthogroups size at a given node in the species tree. Error bars on panels A&C show standard error of the mean value across all true orthogroups.

### Total ancestral genome sizes

We next assessed the accuracy of overall ancestral genome size inference, defined as the total number of genes at each ancestral node in the species tree. For this comparison, we use the same data as the orthogroup size benchmark, we simply sum across all orthogroups to obtain total ancestral genome sizes.

For both the fungi and the plant dataset, ancestral genome sizes inferred by GLADE were significantly correlated with the true ancestral genome sizes (fungi: Spearman’s ρ=0.945, p<2.2×10^-16^; plants: Spearman’s ρ=0.970, p<2.2×10^-16^) (Figure 7). Correlations were also significant for BadiRate (fungi: Spearman’s ρ = 0.975, P<2.2×10^-16^; plants: Spearman’s ρ=0.945, p<2.2×10^-16^) and ancestral character estimation (fungi: Spearman’s ρ=0.968, p<2.2×10^-16^; plants: Spearman’s ρ=0.979, p<2.2×10^-16^) on both datasets, indicating that all three tools reliably recover ancestral genome sizes. The correlations for CAFÉ were much lower due to the significant errors which CAFÉ makes on some nodes (fungi: Spearman’s ρ=0.622, p<2.2×10^-16^; plants: Spearman’s ρ=0.030, p=0.840) (Figure 7A&B).

**Figure 7:**
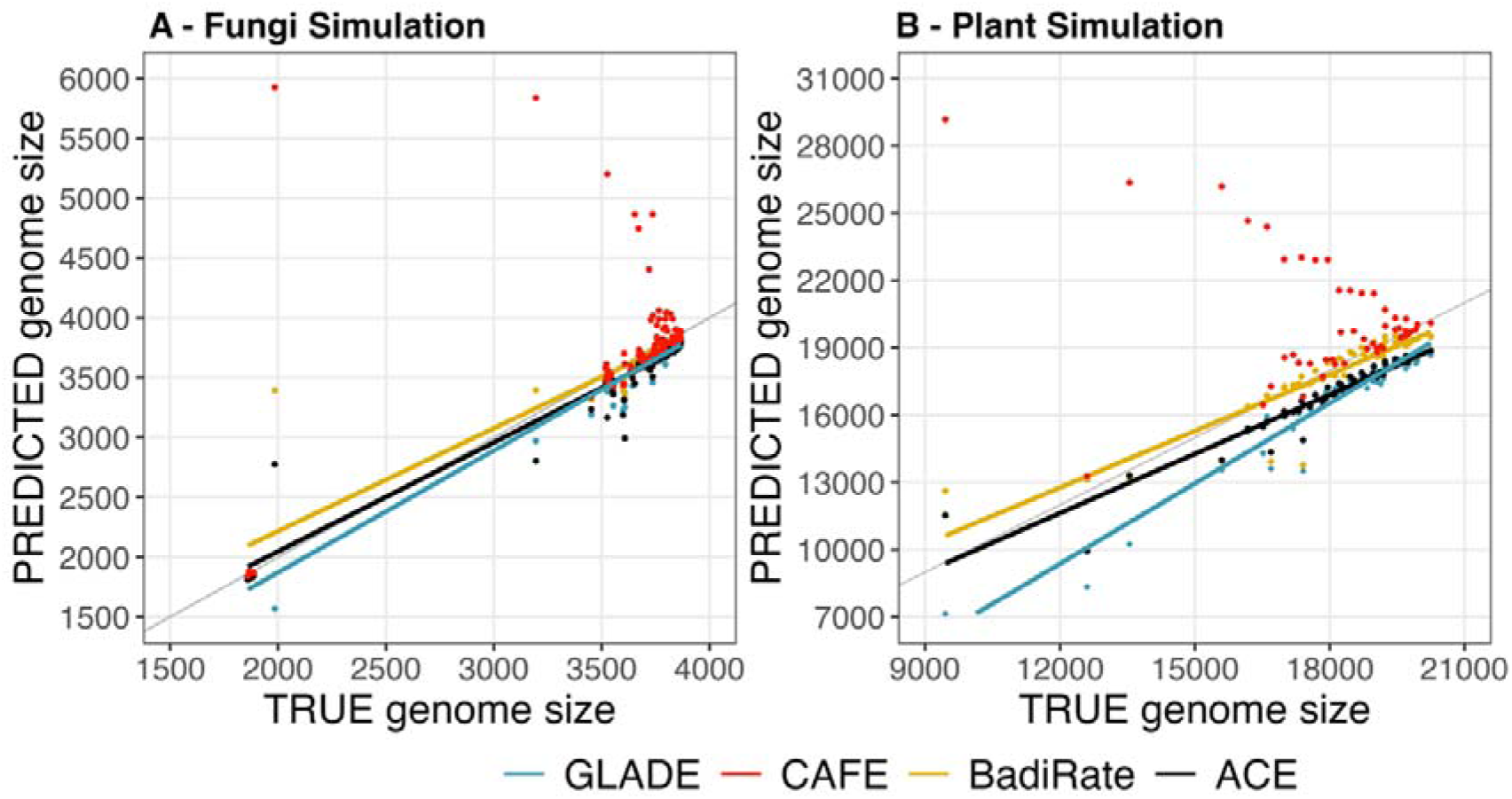
Benchmarking of GLADE on overall genome size (number of genes) for ancestral nodes in the species tree. Grey line shows where true genome size equals predicted genome size.

We further tested whether errors in genome size varied systematically with phylogenetic depth by examining correlations between the magnitude of genome size errors and node position relative to the root. GLADE showed no correlation between node depth and error magnitude in either dataset (fungi: Spearman’s ρ=0.113, p=0.265; plants: Spearman’s ρ=0.067, p=0.645) indicating consistent performance across both shallow and deep nodes. In contrast, ace exhibited a significant but moderate association between error and node depth in both datasets (fungi: Spearman’s ρ=0.461, P<1×10^-5^; plants: Spearman’s ρ=-0.358, p=0.012), and CAFE showed a strong dependence on node depth, with errors increasing markedly toward deeper nodes (fungi: Spearman’s ρ=0.609, p<1×10^-10^; plants: Spearman’s ρ=0.754, p<2.2×10^-16^). Together, these results indicate that methods differ in their sensitivity to phylogenetic depth, with implications for reliability as analyses scale to larger or deeper datasets. Notably, GLADE showed no detectable depth-dependent increase in error in either benchmark.

### GLADE identifies genetic signatures of tooth loss during the evolution of myrmecophagy in mammals

To demonstrate the utility of GLADE, we applied it to investigate extreme dietary specialization in mammals. Some mammals are specialized to feed on ants and termites and eat almost nothing else. This dietary specialization, known as myrmecophagy, has evolved independently at least 12 times across mammals (Vida et al. 2025). Myrmecophagy is phenotypically associated with a loss or simplification of teeth and taste buds, and a lengthening of the tongue and enlarged salivary glands (Reiss 2001; Ferreira-Cardoso et al. 2019). Recent comparative genomic studies have analysed genomes to show the genomic signatures of these patterns, finding that myrmecophagous lineages have frequently lost (through pseudogenization) genes involved in enamel formation and gustation (Emerling et al. 2026). The multiple independent origins of myrmecophagy, the availability of genome data, and the well-characterized phenotypic trait changes make this a good test case to demonstrate the utility of GLADE. Specifically, this enables us to test whether the well-established phenotypic convergence in myrmecophagous mammals is reflected by parallel changes in gene family evolution. For orthogroups associated with tooth formation and taste, we expect to see decreases in orthogroup size at the branches leading to myrmecophagy. For orthogroups associated with salivary gland, we expect to see increases in orthogroups size at the branches leading to myrmecophagy.

We used GLADE to reconstruct gene family evolution across 78 mammalian species, focusing on lineages that independently evolved myrmecophagy (pangolins, echidna, aardvark, and elephant shrew). Orthogroup size changes were inferred by GLADE for all nodes on the species tree, but we focussed on all expansions and contractions along the myrmecophagous branches. Expanded and contracted orthogroups were then functionally annotated using EggNOG-mapper (Cantalapiedra et al. 2021), and Gene Ontology (GO) (The Gene Ontology Consortium 2025) enrichment analyses were performed using ClusterProfiler (Yu et al. 2012) to identify biological processes disproportionately affected by gene family change. This analysis was performed separately for expanded and contracted orthogroups. The most significantly enriched terms were strongly linked to tooth and enamel development, including positive regulation of tooth mineralization, regulation of enamel mineralization, and structural constituent of tooth enamel (Supplementary Figure 1A). These results are consistent with the extensive dental reduction and enamel loss observed in myrmecophagous lineages.

We next examined the gene families inferred by GLADE to have expanded on branches leading to myrmecophagous lineages (Supplementary Figure 1B). Enriched GO terms among these orthogroups were dominated by oxidoreductase activity, alkaloid and icosanoid catabolic processes, and linoleic acid epoxygenase activity, all of which are broadly involved in lipid metabolism and detoxification. These processes may reflect the requirement for myrmecophagous mammals to detoxify various toxic substances (e.g. formic acid and alkaloids) which ants and termites secrete (Cheng et al. 2022).

We then focussed particularly on teeth, as this is the best-studied phenotypic consequence of myrmecophagy. Six orthogroups have enriched functions in tooth formation. Across the phylogeny, GLADE identified 20 instances where the size of one of these tooth-formation orthogroups decreased (Figure 8). Myrmecophagous branches are significantly overrepresented for tooth-associated orthogroups contractions, capturing 40% of tooth-formation orthogroup contractions whilst representing <3% of all branches (binomial test p<1×10^-8^). This provides strong evidence for repeated, convergent contraction of tooth-related orthogroups in ant and termite eating mammals.

**Figure 8:**
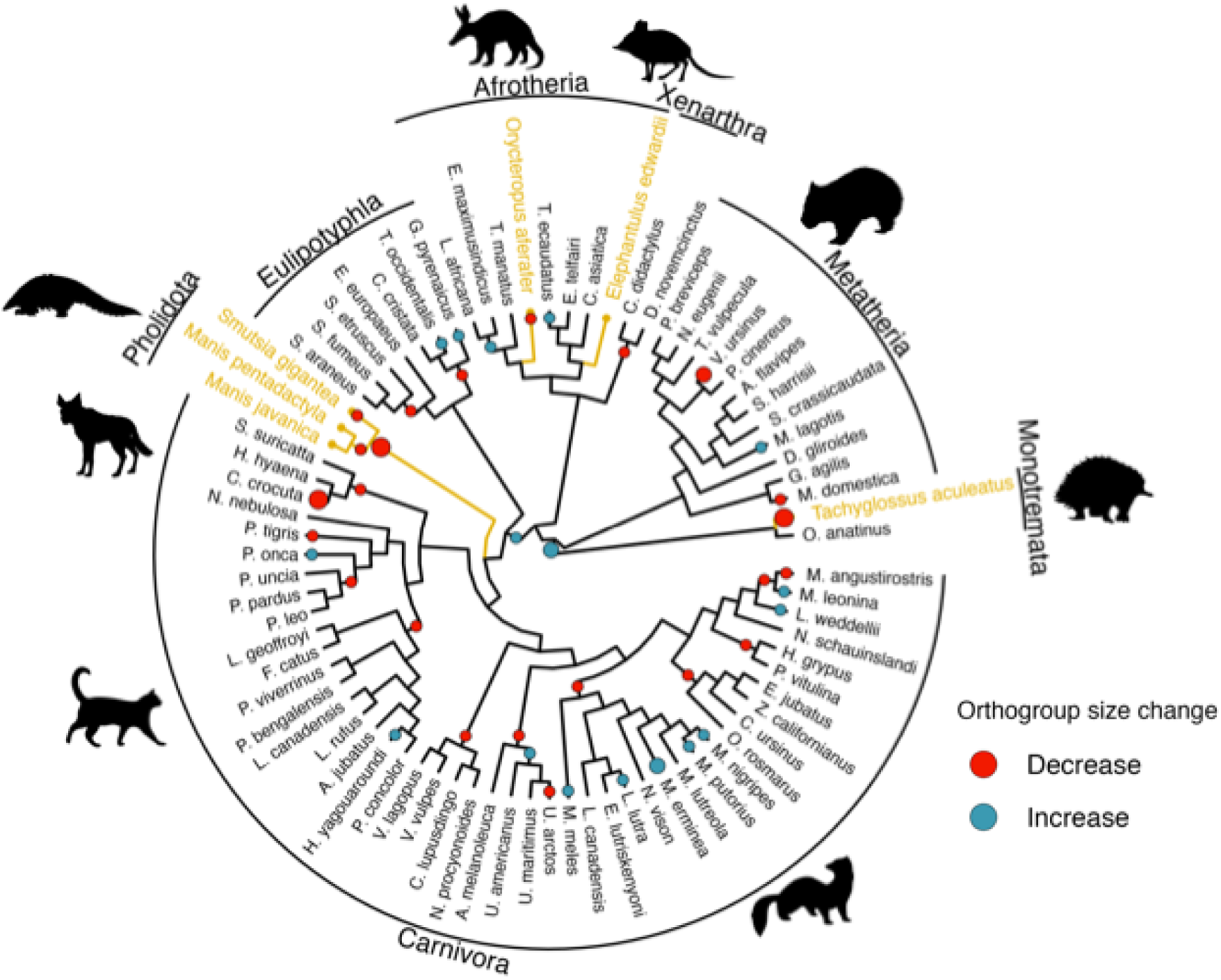
Phylogeny of 78 mammals. Branches and tips in orange show myrmecophagous mammals, whose diet is >90% ants and termites. Circles on nodes show summed changes in orthogroups size for six orthogroups which are enriched for enamel and tooth mineralization. Phylogeny trimmed from (Upham et al. 2019). Silhouettes from phylopic (phylopic.org)

Among non-myrmecophagous species, a small number of contractions were observed on branches leading to the spotted hyena (*Crocuta Crocuta*) and wombat (*Vombatus ursinus*). It is noteworthy that both species have unusual teeth. The spotted hyena is the only carnivorous mammal to be born with a full set of teeth (Pournelle 1965), and the wombat is the only marsupial with continuously growing teeth (Clarke 2003). These contractions may therefore reflect remodelling of the dental toolkit, rather loss *per se*. Together, these results demonstrate that GLADE can detect convergent genomic signatures of adaptation across deep evolutionary time, linking repeated contractions in tooth-associated orthogroups to the emergence of myrmecophagy in mammals.

## Discussion

Expansions and contractions in orthogroups drive evolution, and have punctuated major innovations and transitions across the tree of life. Here, we present GLADE, an easy-to-use tool that facilitates the analysis of these expansions and contractions by mapping gene duplication, gain, and loss over evolutionary time. We demonstrated the accuracy of GLADE on a realistic simulated data and showed that it is substantially more accurate than any competitor method. We further demonstrated the utility of GLADE by analysing a set of 23,000 orthogroups in 78 mammal genomes, identifying convergent contractions in orthogroups linked to tooth formation in ant- and termite eating species. GLADE is presented as general method to enable gene content evolution in the context of adaptation and diversification across the Tree of Life.

To demonstrate GLADE’s accuracy, we applied it to two realistic simulated datasets consisting of sets of genomes where the complete history of every gain, loss, and duplication event in every orthogroup was known (Figure 2). Simulated datasets were necessary here as the true history of each gene family must be known in order to determine whether the events identified are correctly placed on the phylogenetic tree. For real datasets where the truth is not known, it is not possible to evaluate the accuracy of the method. Furthermore, to prevent biasing our method to any particular orthology inference method we trained our simulations on orthogroups inferred by multiple different orthology inference methods. We compared GLADE to FastOMA, as CAFÉ only reconstructs orthogroup sizes rather than directly inferring evolutionary events. GLADE outperformed FastOMA on all event detection metrics on all simulated datasets, achieving a higher accuracy for placement of gene gains in the species tree (F1 scores of 97 & 92 vs. 88 & 72: Figure 3), a higher accuracy detecting the branches where gene losses occurred (F1 scores of 81 & 63 vs. 68 & 64: Figure 4), and a higher accuracy in identifying and locating gene duplication events in the species tree (F1 scores of 87 & 81 vs. 70 & 55: Figure 5). Where GLADE made errors they tended to be small, and for losses were significantly smaller on average than errors made by FastOMA (Figure 4).

Where GLADE and CAFÉ could be compared, GLADE outperformed CAFÉ at reconstructing ancestral orthogroup sizes (Figure 6). Across both fungal and plant simulations, GLADE achieved the highest overall accuracy, as reflected by its F1 scores (fungi: 94.9; plants: 86.8). Both CAFÉ and BadiRate exhibited recall values that were comparable to or slightly higher than those of GLADE (fungi: CAFÉ = 98.9, BadiRate = 98.4, GLADE = 98.6; plants: CAFÉ = 93.1, BadiRate = 90.9, GLADE = 92.5). However, this was offset by substantially lower precision for both methods relative to GLADE (fungi: CAFÉ = 86.3, BadiRate = 84.9, GLADE = 91.4; plants: CAFÉ = 70.3, BadiRate = 72.2, GLADE = 81.8), resulting in lower overall accuracy. CAFÉ has a tendency to infer events to have occurred on deeper branches in the species tree than GLADE, due to difficulties of maximum likelihood approaches in successfully detecting convergent correlated changes in neighbouring branches. Errors in species tree inference and assignment of genes to orthogroups will likely propagate to errors in gene events, but both CAFÉ and GLADE require users to provide this information, and many users first run OrthoFinder to obtain the required input for CAFÉ. With efficient parallelization, GLADE is also more easily scalable to large analyses than CAFÉ, and provides a richer output including ancestral genomes at every node, containing the relevant number of representative sequences from each orthogroup.

Across both fungal and plant benchmarks, methods differed in their robustness to phylogenetic depth when reconstructing total ancestral genome sizes (Figure 7). Although GLADE, BadiRate, and ancestral character estimation all showed strong correspondence with true ancestral genome sizes (Spearman’s ρ > 0.94 in both datasets), only GLADE showed no detectable increase in error toward deeper nodes (fungi: ρ = 0.113, plants: ρ = 0.067: Figure 7C&D). In contrast, ACE and especially CAFÉ exhibited significant depth-dependent error, with CAFÉ showing strong positive correlations between error and node depth (fungi: ρ = 0.609; plants: ρ = 0.754). Together, these results indicate that while several methods recover relative ancestral genome size trends, GLADE provides more stable inference across both shallow and deep nodes, which is likely to improve reliability in large or deeply sampled phylogenies. To quantify absolute reconstruction accuracy, we additionally calculated accuracy as one minus the root mean squared error (RMSE) between reconstructed and true ancestral genome sizes across all nodes (Figure 7E&F). In the fungal simulations, GLADE scored the highest accuracy (0.952), outperforming ancestral character estimation (0.945) and BadiRate (0.927), while CAFÉ showed substantially lower accuracy (0.770). In the plant simulations, BadiRate achieved the highest accuracy (0.934), followed by ancestral character estimation (0.923), whereas GLADE exhibited slightly lower accuracy (0.895) and CAFÉ again performed worst (0.623). Taken together, our results show that GLADE offers a good balance between overall accuracy and robustness to phylogenetic depth.

To demonstrate the real-world utility of GLADE, we applied it to investigate extreme dietary specialization in mammals (Figure 8). Myrmecophagy – a diet of over 90% ants and termites – has evolved at least 12 times within mammals, across taxonomically distinct lineages such as pangolins, aardvarks, anteaters, and armadillos (Vida et al. 2025). This adaptation is linked to the expansion in social insect abundance during the Cenozoic, which is itself linked to the diversification and spread of angiospems, particularly into open habitats (Nelsen et al. 2023). Loss or reduction in teeth is a well-known characteristic of myrmecophagy, but whether the convergent loss of teeth is associated with convergent changes in gene content is less well understood. However, a recent innovative recent study (Emerling et al. 2026) found that certain genes involved in enamel formation have become convergently pseudogenised in almost all myrmecophages for which genome data was available. To demonstrate the utility of GLADE In our analysis using GLADE, we showed that gene functions linked to tooth and enamel development are strongly enriched in orthogroups that decrease in size on branches where myrmecophagy evolved. Looking specifically at six key tooth-formation orthogroups, we showed myrmecophagous lineages are significantly overrepresented in cases where one of the orthogroups decreased in size along a branch of the phylogeny (40% of 20 instances, Figure 8). We focused on teeth and dentition so that we could compare to known phenotypic changes, but GLADE could also provide unexpected findings on other orthogroup changes that occur repeatedly on myrmecophagy branches.

Beyond the analyses presented here, GLADE enables a wide range of evolutionary investigations. GLADE returns ancestral gene repertoires alongside the full set of mapped events, allowing it to be integrated with complementary approaches to link genomic change to phenotypic innovation (Pagel 1994; Cornwallis and Griffin 2024; Dewar et al. 2025). For example, ancestral reconstructions of ecology or behaviour can be mapped onto the same phylogeny, allowing researchers to test whether particular genomic events preceded, coincided with, or followed key evolutionary transitions (Feng et al. 2024). Combining GLADE with functional annotation tools also enables hypotheses about how molecular functions expanded, contracted, or diversified through time – such as exploring the evolution of orthogroups underlying cooperation or diet (Hao et al. 2024; Hoile et al. 2025). GLADE’s reconstructions can also be paired with analyses of molecular evolution to identify where changes in selective pressures coincide with orthogroup expansions and contractions. These potential integrations make GLADE a powerful and flexible platform for studying genome evolution.

In conclusion, GLADE is a valuable tool for reconstructing the evolutionary history of orthogroups. GLADE is easy to use, requiring only an OrthoFinder results directory as input and no user specified parameters. The output of GLADE includes the complete identification of gain, loss, and duplication events in the history of every orthogroup, as well as reconstructed ancestral genomes at every internal node in the species tree. GLADE is available on GitHub at https://github.com/lauriebelch/GLADE/.

## Methods

### Implementation

GLADE is a python application designed for orthogroup evolutionary analysis through the identification of gene gain, loss, and duplication events across the evolutionary history of each orthogroup. It requires Python >=3.9 and integrates several python libraries, including ETE3 (Huerta-Cepas et al. 2016) for tree parsing and traversal, and numpy for numerical computation. GLADE uses a modular architecture to perform its orthogroups reconstruction. These modules work in concert to infer ancestral gene content, map orthogroup changes across branches, and compute duplication, gain, and loss statistics. Performance is enhanced through the use of multiprocessing, enabling parallel computation of branch-specific or orthogroups-specific events. GLADE is compatible with macOS, Linux, and Windows systems, and can be run using a single python script. Comprehensive installation and usage instructions for GLADE are available at https://github.com/lauriebelch/GLADE.

### GLADE workflow

#### Input Files

GLADE requires only a complete OrthoFinder v3 (Emms et al. 2025) run as input. The specific files that are used are 1) Mapping of genes to orthogroups (Orthogroups/Orthogroups.tsv), 2) Gene trees for each orthogroups (Resolved_Gene_Trees/Resolved_Gene_Trees.txt), 3) A species tree (Species_Tree/SpeciesTree_rooted_node_labels.txt). This can be either a tree that users have supplied to OrthoFinder, or the one generated by the OrthoFinder algorithm.

#### Orthogroup filtering

Before analysis, orthogroups containing fewer than four total gene copies across all sampled taxa are removed. This step eliminates very small groups for which many tree inference tools won’t generate trees, and which can lead to unreliable event inference. Remaining orthogroups are processed in downstream analyses.

#### Orthogroup gain identification

An orthogroup can exist at the root of the species tree, or might emerge later on. GLADE infers orthogroups gain by traversing the species tree from root to tip. For any node, there are two descendant clades (Child A and Child B), each of which have genes from certain species (Species set A and Species set B). An orthogroup is considered present at a node if it has genes from both species sets. The node at which the orthogroups was gained is the most-ancestral node at which the orthogroups is present. This is equivalent to the phylostratigraphy approach (Domazet-Lošo et al. 2007). Gains are annotated to the branch connecting this node to its parent.

#### Gene duplication events

GLADE infers duplications in an orthogroup by traversing the gene tree from root to tip, using the same method as STRIDE (Emms and Kelly 2017). For each node there are two descendant clades (Child A and Child B), each of which have genes from certain species (Species set A and Species set B). If there is overlap in the two species sets, GLADE infers a duplication at that node. Each duplication is given a support value defined as the proportion of expected species shared by both child clades. Duplications with support ≥0.5 are retained and labelled as high-confidence events. Lower-support duplications are recorded separately but not counted in branch-level aggregation.

#### Loss

GLADE infers two types of gene loss. One type is ‘speciation loss’, which occur when a species which emerged through speciation after the orthogroup itself emerged has no gene in that orthogroup. The other type is ‘post-duplication loss’, which occur when a species doesn’t retain all expected copies after a gene duplication event.

#### Speciation loss

To identify speciation loss in an orthogroups, the species tree is traversed from the inferred gain node. For every internal node, the two descendant clades (Child A and Child B) are examined for genes from that orthogroup. If one clade contains at least one member of the orthogroup while its sister clade contains none, GLADE infers a loss along the branch leading to the lineage that lacks the orthogroup. Each event is recorded with the node name, relevant species, and orthogroup identity.

#### Post-duplication loss

To identify post-duplication losses in an orthogroup, we traverse the gene tree from root to tip. For every high-confidence gene duplication node in a gene tree, the two descendant clades (Child A and Child B) are identified. The species represented in each clade (Species set A and Species set B) are compared to the set of species expected based on the corresponding node in the species tree. When a species is present in one child clade but missing in the other, GLADE infers that the lineage has lost one of the duplicated copies. Each loss is linked to its originating duplication event and annotated with the relevant species and node identifiers.

#### Mapping events to branches

GLADE maps each inferred gain, loss, and duplication event onto a specific branch of the species tree to quantify branch-wise evolutionary dynamics. The species tree is traversed to define all nodes and there parent-child relationships. For each orthogroup, the gain node, speciation losses, post-duplication losses, and duplication events are assigned to their corresponding branches based on these node relationships. Gains are mapped to the branch connecting the parent of the gain node to the gain node itself, whereas speciation losses are mapped to the branch connecting the parent node to the child clade that lacks the orthogroup. Duplications are assigned to the branch leading to the duplication node in the species tree, and post-duplication losses are mapped to the branch where a descendant species fails to retain one of the duplicated copies. The resulting counts of gains, losses, and duplications per branch are summarized in a single output file (Branch_statistics.tsv), with separate files listing the orthogroups involved with each event type.

#### Ancestral genome reconstruction

Ancestral gene content for each internal node in the species tree is inferred for every orthogroup. An orthogroups ancestral count is zero for all internal nodes prior to the node at which the orthogroups emerged. For all other species tree nodes, orthogroup membership at that node is reconstructed using the gene tree. Let us consider species tree ancestral node N1, and orthogroup OG002. We first define the set of descendant species under node N1. We then prune the gene tree to include only genes from those species. Some of these genes might be the result of duplications that occurred later in the species tree than N1. We use our mapping of duplications to the species tree to identify these events, and randomly remove one of the descendant branches. This makes the gene tree representative of the reconstructed situation at the time of N1.

Duplications that occurred before the focal node are summed to estimate the number of expected ancestral copies. To select which sequences to include in the ancestral genome, GLADE first calculates root to tip distance for all leaves in the pruned gene tree. Leaves are then iteratively sampled from the gene tree, with the leaf closest to the median root to tip distance sampled each time. This continues until either the number of selected leaves matches the number of expected ancestral copies for that orthogroups, or until there are no more leaves in the gene tree. If fewer sequences are available than the expected number of ancestral copies, GLADE treats this as implied gene losses along the descendant lineages. The process is repeated for all orthogroups and all nodes, and the resulting ancestral gene set is compiled into a FASTA file for each internal node, representing the inferred ancestral genome. Orthogroup sizes for both extant and ancestral species are quantified by counting the number of genes belonging to each orthogroup in these reconstructed genomes. It should be noted that our approach attempts to sample those sequences that have experienced least sequence change since divergence from the ancestral node under consideration. Thus, they are not reconstructed ancestral sequences (and should not be treated as such), and are instead provided to enable users to make use of these reference sets for constructing databases or performing BLAST searches for phylogenetic profiling or similar analyses.

#### Quantifying orthogroups size changes on branches

GLADE quantifies changes in orthogroup size along each branch of the species tree by integrating data from the orthogroup counts in extant species and the orthogroups counts inferred in ancestral nodes. For every orthogroup, the number of genes present in each ancestral and extant species is obtained from the reconstructed genomes. For each branch, GLADE retrieves the orthogroup size at both the parent and the descendant node. The difference between these values is the inferred change in orthogroup size along that branch.

As an additional feature, GLADE also provides change normalised by branch length. Branch lengths are obtained from the species tree, and the absolute value of each orthogroup’s size change is divided by the relevant branch length to obtain an estimate of rate of change per unit of evolutionary time.

#### Output Files

GLADE generates a set of output files summarizing orthogroup evolution across the species tree. The GainsLossDuplication output directory contains files listing all identified events across all orthogroups, with all nodes mapped to their position on the species tree.

GainsLossDuplication/Gains.tsv, Loss_speciation.tsv, Loss_postduplication.tsv, and Duplications.tsv list, for each orthogroup, the nodes or branches where gene gains, losses, or duplications occurred. Branch_statistics.tsv summarizes the total number of each event type per branch, while the corresponding *_bybranch.tsv files provide the expanded lists of orthogroups associated with each branch-level event.

Ancestral gene repertoires are stored in AncestralGenomes/, with one FASTA file per internal node representing the reconstructed ancestral genome. AncestralGenomes.txt provides summary statistics (gene counts and total sequence length), and Ancestral_HOG_counts.csv records orthogroup copy numbers for all extant and ancestral species.

For extant taxa, orthogroup counts are listed in GainsLossDuplication/extant_OG_counts.tsv. Finally, OrthogroupBranchChange.tsv integrates these datasets to quantify gene family size change along each branch, reporting the number of genes at parent and descendant nodes, branch length, net change, and rate of change per unit branch length.

### Orthogroup simulations

Methods for orthogroup simulations are provided in Supplemental File 1. All simulated genomes and results files are available at 10.6084/m9.figshare.31158355.

### Benchmarking

Methods for benchmarking are provided in Supplemental File 2.

### Application of GLADE to mammalian genomes

#### Selection and preparation of Genomes

We used mammalian genomes from NCBI that have proteomes. We filtered to include only genomes from the section of the Upham et al. mammalian phylogeny (Upham et al. 2019) that has most ant-eating species (*Monotremata, Metatheria, Afrotheria, Xenarthra, Eulipotyphla, Pholidota, Carnivora*). We used NCBI Datasets (O’Leary et al. 2024) to download the proteomes, and OrthoFinder’s primary transcripts script to only include the longest isoform for each gene. This dataset has 6 ant-eating species (Giant Pangolin *Smutsia gigantea*, Chinese Panoglin *Manis pentadactyla*, Sunda Pangolin *Manis javanica*, Short-beaked echidna *Tachyglossus aculeatus,* Aardvark *Orycteropus aferafer*, and Cape elephant shrew *Elephantulus edwardii*)

#### Phylogeny

We used the phylogeny from Upham et al. (2019). Myremcophagy has evolved in 4 branches on this tree: the branch leading to *Manidae* (N10-N16), the branch leading to *Tachyglossidae* (N1-*Tachyglossus aculeatus*), the branch leading to *Orycteropodidae* (N19-*Orycteropus aferafer*), and the branch leading to *Macroscelididae* (N18-*Elephantulus edwardii*). We used these branches as our focal myrmecophagy branches.

#### Analysis of orthogroup dynamics

We ran OrthoFinder v3.1 and GLADE v1 on the 78 proteomes with default settings. We used the orthogroups size changes inferred by GLADE, and filtered to include only the focal myrmecophagy branches. Orthogroups showing a decrease in size on the focal ranches were classified as contracted, whereas orthogroups showing an increase in size were classified as expanded. For each contracted or expanded orthogroups, we retrieved all of its member genes from the OrthoFinder output. This resulted in two gene sets: genes from contracted orthogroups, and genes from expanded orthogroups.

We obtained functional annotations from all genes across all species using EggNOG-mapper (Cantalapiedra et al. 2021). We retained all unique annotated genes to define the ‘background’ genes for further analysis. We then used the R package ClusterProfiler (Yu et al. 2012) to perform Gene Ontology (GO) enrichment analysis. We performed separate test for the expanded and contracted genes compared to the background genes, using the Benjamin–Hochberg correction p-value correction for multiple testing. To identify broad gene functions, we retrieved GO term names using the R package ontologyIndex (Greene et al. 2017), and computed semantic similarity between enriched GO terms using the Wang method implemented in the R package GOSemSim (Yu et al. 2010). Redundant terms were then collapsed using ClusterProfiler’s simplify function. We then used eggnog-mapper to functionally annotate all genes across the 78 proteomes.

From the GLADE output, we extracted a list of orthogroup+branch combinations where the orthogroups changes in size on a myrmecophagy branch. We then extracted the genes in these orthogroups, giving a list of genes from orthogroups that decrease in size in focal branches (loss genes), and a list of genes from orthogroups that increase in size in focal branches (gain genes). Next, we used the R package ClusterProfiler to look at functions that are enriched in each of these sets, compared to a background of all genes. We calculated GO semantic similarity scores using the Wang Method, using a semantic similarity cutoff of 0.65 to retain representative GO terms.

## Acknowledgements

This work was funded by the Wellcome Trust under grant agreement number 226598/Z/22/Z.

