## Supplemental File 1 for "GLADE: Accurate inference of Gains, Losses, Ancestral genomes, and Duplication Events for comparative genomics"

Orthogroup simulations

To benchmark GLADE, we used simulated orthogroups. This approach allows us to know exactly which gene belongs to which orthogroup and exactly where each gene gain, loss, and duplication occurred within the evolutionary history of that orthogroup. We produced two simulated datasets. One for fungi, and one for plants. For the fungi simulation, we obtained N=100 species of budding yeast from the Shen et al. (2018) budding yeast data by subsampling 100 species using Phylogenetic Diversity Analyzer (PDA) (Chernomor et al. 2015) from the larger tree. This provided us with a set of 100 proteomes and a corresponding species tree. For the fungi simulation, we used *Saccharomyces cerevisiae* as the starting genome. For the plant simulation, we obtained N=50 species of plants from the Ensembl plants dataset (Bolser et al. 2016) by subsampling 50 species using PDA. For the plant simulation, we used *Arabidopsis thaliana* as the starting genome.

For both simulations, our approach involves picking a genome to designate as the ancestral genome. We then simulate evolution of each gene in that genome along a species tree. In this way, we are matching the phylogenetic definition of an orthogroup as the set of genes descended from a single gene in their most recent common ancestor. In order to create realistic simulations, we use real data for these two features. We use a gene from the real genome (*S. cerevisiae* for fungi or *A. thaliana* for plants) to seed each orthogroup, and use the relevant phylogeny (N=100 fungi or N=50 plants) as the species tree. We also use empirically estimated parameters to inform how we simulate evolution in each orthogroup (see section on ‘parameter estimation’). We also ran PfamScan (Finn et al. 2014) on the starting genome to identify domains. This allows us to inform our evolutionary simulations about which regions of a protein should be more or less conserved during the simulated evolution.

For each simulated orthogroups, we follow the procedure outlined below. First, we randomly select a gene from the starting genome. Each gene can only be used in one simulated orthogroup. Next, we simulate a gene tree using SagePhy (Kundu and Bansal 2019) GuestTreeGen. These simulations are given a value for duplication rate, loss rate, and gene-birth-coefficient (representing how biased orthogroup birth is towards the root of the species tree). See the section on ‘parameter estimation’ to see how these values are determined. At this stage, a simulated gene tree will be ultrametric (all leaves are equidistant to the root). We therefore use SagePhy BranchRelaxer to relax the branch lengths to provide realistic branch lengths. We use the ACRY07 relaxation model which requires a start rate and variance. As empirical estimates were not available for these parameters, we set a maximum start rate of 0.5 and a maximum variance of 0.1, from which a value for a given orthogroups was randomly drawn. Next, we simulate a multiple sequence alignment for the relaxed gene tree. If the gene tree doesn’t begin at the root of the species tree, we shuffle the sequence before starting this step to represent the birth of a novel gene. We used IQ-TREE Alisim (Ly-Trong et al. 2022; Wong et al. 2025) to simulate alignments, as this allows us to define partition files providing different parameters for different regions of our starting protein. Domain regions are defined as any region flagged as a domain or motif by PfamScan, and Non-domain regions are all other regions. Domain regions were given an amino acid substitution model generated from our empirical parameter estimation. Non-domain regions were given a JTT+G4+I substitution model, with gamma shape alpha and proportion of invariant sites informed by our empirical parameter estimation. All regions were given indels (0.04 for max insertion or deletion rate, and 1 for size).

For the fungi simulation, we obtained a total of n=5917 orthogroups. For the plant simulation, we obtained a total of n=27000 orthogroups. To turn our simulated alignments into genomes, we extracted sequences and saved them into a different proteome file for each species.

Simulation parameter estimation

To inform our simulations and ensure that they reflect realistic patterns of molecular evolution, we estimated certain parameters from empirical data. We did this separately for the fungi and the plant simulations. For each simulation, we used the real genomes to estimate substitution model parameters and duplication/loss rates. Orthogroups were identified using OrthoFinder, and protein sequences for each orthogroup were then aligned with MAFFT (Katoh and Standley 2013) --localpair option and trimmed with trimAl (Capella-Gutiérrez et al. 2009) -automated1. We then used IQTree3 to estimate substitution model parameters. We did this by running IQTree3 ModelFinder (Kalyaanamoorthy et al. 2017) for JTT+G4+I models. From each analysis, we extracted the gamma shape parameter (alpha) and the proportion of invariant sites (I) under the best-fitting model. Across all orthogroups, the mean and standard deviation of alpha and I were calculated, and these values were used to parameterize log-normal distributions from which simulation values were drawn.

To approximate realistic rates of gene duplication and loss, we used OrthoFinder to count the average number of duplication events per orthogroup. This count forms are average duplication rate to be fed into gene tree simulation. Because gene loss rates cannot be directly estimated from these data, we set the average loss rate to be 1.5 times higher than the duplication rate, consistent with empirical evidence that losses are more common than duplications (Birchler and Yang 2022).

To assign substitution models to protein domains, we used the PfamScan output for the *S. cerevisiae* genome to identify domain regions. For each domain region, we used IQTree3 ModelFinder to find the best fitting amino acid substitution model. These models were stored as a list, to allow them to be randomly sampled during simulations.

Simulation validation

To assess whether the simulated orthogroups resembled real orthogroups in their basic properties and tree structures, we compared several statistics between simulated and empirical datasets. For the empirical reference, we analyzed orthogroups inferred from the real genomes using OrthoFinder (Emms et al. 2025), Broccoli (Derelle et al. 2020), SonicParanoid2 (Cosentino et al. 2024), and FastOMA (Majidian et al. 2025), four widely used and independently developed orthology inference tools. For both simulated and empirical datasets, we summarized the distribution of orthogroup sizes in terms of the number of genes per orthogroup and the number of species represented.

To evaluate the realism of the simulated gene trees, we used two measures. The Wiener index is the sum of the shortest path distances between all pairs of nodes in a tree, providing a measure of tree shape and balance (Chindelevitch et al. 2021). The ‘treeness’ index is the proportion of total branch length that is contained in internal (nor-terminal) branches, reflecting the extent to which evolutionary change is concentrated in leaf versus internal branches (Astolfi and Zonta-Sgaramella 1984). We performed these analyses using the R package treestats (Janzen and Etienne 2024).
