## Supplemental Figures for "GLADE: Accurate inference of Gains, Losses, Ancestral genomes, and Duplication Events for comparative genomics"

**Supplementary Figures**

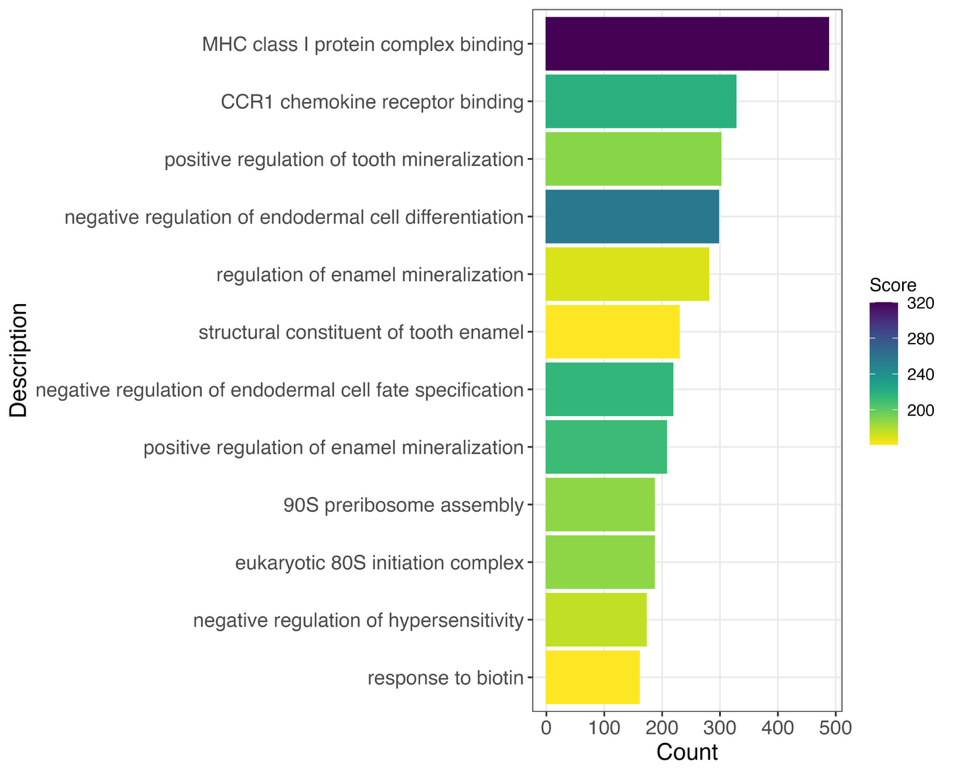

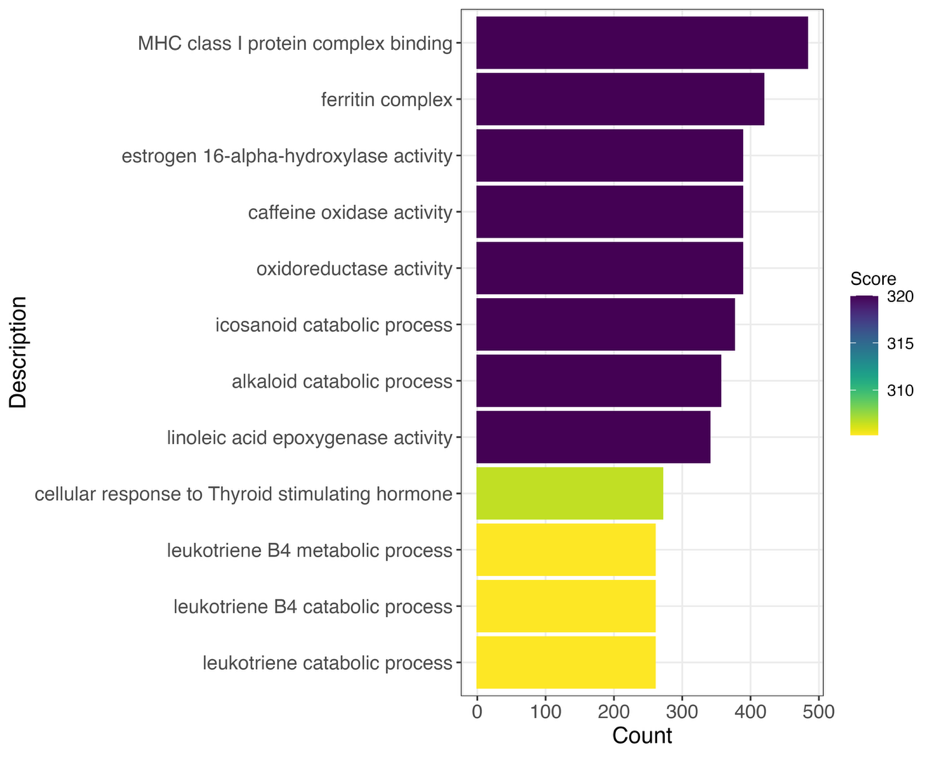

**Supplementary Figure 1**: GO enrichment analysis for orthogroups that change size on branches leading to myrmecophagy. Top panel shows orthogroups that decrease in size on these branches, bottom panel shows orthogroups that increase in size on these branches. X-axis shows count, and colour shows score, calculated as -log10 of adjusted p-value.
