## Supplemental File 2 for "GLADE: Accurate inference of Gains, Losses, Ancestral genomes, and Duplication Events for comparative genomics"

Benchmarking

Benchmarking ancestral genomes

To benchmark the ancestral orthogroup sizes inferred by GLADE, we compared its results to three alternative methods for reconstructing gene family evolution – CAFÉ (Mendes et al. 2021), BadiRate (Librado et al. 2012), and ancestral character estimation (ace) (Revell 2014) implemented in the R package ape (Paradis and Schliep 2019). All methods require data on orthogroup counts in extant species, which we generated for all methods using OrthoFinder, so that any differences between the two methods are due to the method for inferring ancestral sizes and not the underlying orthology inference method. We ran CAFÉ with among-family rate variation and three discrete gamma rate categories (using the -k option) and the -z option to infer ancestral sizes for orthogroups that don’t exist at the root. We ran BadiRate with default parameters. We ran ancestral character estimation with an ‘equal rates’ model and discrete states each representing an orthogroup size (e.g. 2).

Ground truth for ancestral orthogroup sizes

To reconstruct the true orthogroup size at every ancestral node in the species tree, we use the intermediate files generated by SagePhy during the gene tree simulation. Specifically, the unpruned.guest2host file contains information on every event (duplication, loss, speciation), mapped to its time and location on the species tree. For nodes prior to the emergence of the orthogroup, the reconstructed size is 0. For subsequent nodes, we add duplications and subtract losses to reconstruct the true orthogroup size.

Measuring precision, recall, and F1 score

For each simulation, there are N simulated orthogroups (5917 for fungi, 27000 for plants), which we term RefOGs. All tools have the same set of inferred orthogroups, which are termed PredOGs. Each simulated gene is associated with a ground-truth ancestral size at all internal nodes. If RefOGs and PredOGs perfectly overlapped, then a tool scores a true positive (TP) if the gene is in an orthogroup with the correct reconstructed ancestral size, a false positive (FP) if the gene is in an orthogroup with incorrect ancestral size, and a false negative (FN) if the gene doesn’t have a predicted ancestral size. However in reality, each RefOG may be split across several PredOGs, or fused with a different RefOG, or either contain extra genes or miss genes. Our aim is to assess how accurate the PredOGs of a tool are in terms of ancestral reconstruction. Therefore, we use a weighting approach to account for imperfect overlap between RefOGs and PredOGs. Specifically, we use the following approach for each RefOG. 1) Identify all overlapping PredOGs which contain at least 1 gene from the RefOG. Each of these PredOGs have a reconstructed ancestral size for every internal node in the species tree. 2) Analyse each PredOG in turn. For each node, a true positive (TP) occurs when a PredOG’s predicted family size matches the true ancestral size for that RefOG. A false positive (FP) occurs when a gene in a PredOG has an incorrect predicted size. A false negative (FN) occurs for genes in the RefOG that are missing from the PredOG (and therefore have no predicted value). 3) Calculate precision (TP / (TP + FP) and recall (TP / (TP + FN) for each PredOG. 4) Calculate an overall recall and precision for that RefOG, by weighting each PredOG score by the proportion of the RefOG genes that it contains. To obtain single measures for precision and recall across all RefOGs, we weight the per-RefOG scores by the size of the RefOG. We then calculate the F1 score as the harmonic mean of precision and recall. We also calculate standard errors of precision, recall, and F1 scores as a measure of variability. Our approach allows us to evaluate how well each method reconstructs ancestral gene family sizes, even when orthogroup boundaries are imperfectly inferred. For each tool we also measured the magnitude of each error made, defined as the absolute difference between true and inferred orthogroup size at a node.

Benchmarking Duplications, Losses, and Gains

Duplications, losses, and gains aren’t directly inferred by CAFÉ, BadiRate, or ace - therefore we benchmark GLADE against FastOMA, using pyHam to extract the relevant evolutionary events. We ran FastOMA on the simulated genomes to generate inferred orthogroups, and then used the pyHam tool from the same research group to extract information on gains, losses, and duplications.

Benchmarking Duplications

To build the ground truth, we used the pruned.guest2host files from SagePhy to extract all of the duplications in the simulations. We then used the gene trees to extract the two set of leaves produced by each duplication (one from each descendant lineage). For each duplication, this leaf set forms our ground truth. For benchmarking, a true positive (TP) occurs when a tool infers a duplication leaf set that is identical to the ground truth. A false positive (FP) occurs when a tool infers a duplication leaf set that doesn’t exist in the ground truth. A false negative (FN) occurs when a true duplication leaf set isn’t inferred by a tool. To calculate the errors made by each tool, we first match each false positive duplication set to its closest true duplication set, using Jaccard index. We then quantify effort as the amount of effort required to turn the false set into a true set, where adding or removing one gene is equivalent to an effort of one.

Benchmarking Losses

To build the ground truth, we use the unpruned.guest2host file from SagePhy to extract nodes on the species tree where loss events have occurred in an orthogroup.

Loss inference accuracy was assessed by comparing inferred loss events to simulated ground-truth losses based on their placement on the species tree. A true positive (TP) was defined as a loss inferred at the same species-tree node as a simulated loss, a false positive (FP) as a loss inferred at a node without a corresponding simulated loss, and a false negative (FN) as a simulated loss for which no loss was inferred at the corresponding node. We use the same weighting approach as is used for ancestral reconstruction benchmarking to calculate overall precision, recall, and F1-score

To quantify errors, we measured the distance between an inferred loss node for a given orthogroups and the nearest true loss node for that orthogroups (if applicable). We then measured that distance in two ways – either topological distance (number of intervening nodes), or branch length distance. For example, if a loss is incorrectly inferred at node N2 and the closest true loss is N4, we could calculate both the shortest topological distance between those nodes (e.g. 2 nodes), and also the branch length on the shortest path between those nodes (e.g. 0.1).

Benchmarking Gains

To build the ground truth, we use the unpruned.guest2host file from SagePhy to extract the node at which an orthogroup first appeared. Each RefOG has a single value for this node.

Gain inference accuracy was assessed by comparing the inferred gain (birth) node of each orthogroup to the simulated ground-truth birth node on the species tree. For overlapping predicted orthogroups, we defined true positives (TP) as recovered genes whose inferred gain node matched the true birth node, false positives (FP) as recovered genes assigned a gain node different from the true birth node, and false negatives (FN) as true orthogroup genes missing from the predicted orthogroups. We use the same weighting approach as is used for loss benchmarking.

Errors were quantified in the same way as they were for gene loss events, using both topological and branch length measures of error magnitudes.
